# Composite plants for a composite plant: An efficient protocol for root studies in the sunflower using composite plants approach

**DOI:** 10.1101/712760

**Authors:** Tyler Parks, Yordan S. Yordanov

## Abstract

Sunflower (*Helianthus annuus L*.) is important oilseed crop in the world and the sunflower oil is prized for its’ exceptional quality and flavor. The recent availability of the sunflower genome can allow genome-wide characterization of genes and gene families. With plant transformation usually being the rate limiting step for gene functional studies of sunflower, composite plants can alleviate this bottleneck. Composite plants, produced using *Agrobacterium rhizogenes*, are plants with transgenic roots and wild type shoots. Composite plants offer benefits over creating fully transgenic plants, namely time and cost. Here we outlined the critical steps and parameters for a protocol for the sunflower composite plants production. We use more than a dozen genotypes and three constitutive promoters to validates the utility and efficiency of this protocol. Moreover, functional gene characterization by overexpression and RNAi silencing of a root related transcription factor HaLBD16 further emphasize the value of the system in the sunflower studies. With the protocol developed here an experiment can be carried out with efficiency and in only two months. This procedure adds to arsenal of approaches for the functional genetics/genomics in sunflower for characterization candidate genes involved in root development and stress adaptation.

**Key message:** Composite plants technique described here is fast and efficient approach for roots functional studies in sunflower.

## MATERIALS AND METHODS

### Consumables and chemicals

Chemicals used were purchased through Sigma-Aldrich (St. Louis, MO) or Fisher Scientific (Pittsburgh, PA). Reagents for GUS staining and antibiotics were purchased from GoldBio (Olivette, MO). Caisson boxes were ordered directly from Caisson Labs (Smithfield, UT). Rockwool was purchased from PowerGrow Systems (Vineyard, UT). All water used in experiments was purified with the Milli-Q Reagent System (Millipore, Billerica, MA).

### Plant Material

Sunflower seeds were ordered mostly from the USDA National Plant Germplasm System and planted in garden beds in the greenhouse courtyard at Eastern Illinois University in May. Variety used: Peredovik (NPGS accession numbers-PI 650338), HA115 (PI 650577), HA236 (PI 650592), HA89 (PI 599773), RHA801 (PI 599768), HIR34 (PI 650613), RHA311 (PI 599789), RHA271 (PI 599786), RHA298 (PI 599766), HA412 (PI 603993), HA412-HO (PI 642777), RHA280 (PI 552943), Mammoth (Garden store). The inflorescences were isolate during pollination to maintain pure lines in self-pollinated varieties. In open-pollinated variety Peredovik four inflorescence of the same variety were uncovered and cross-pollinated by hand to obtain seeds. Seeds were harvested in mid-September 2016, cleaned, and stored at 4**°**C.

### *Agrobacterium* strain and plasmid constructs

*Agrobacterium rhizogenes* strain K599 (Mankin et al. 2007) was used to infect the hypocotyls of sunflowers. Plasmids with the desired insert were transformed into electrocompetent *A. rhizogenes* strain K599 using the Bio-Rad Gene Pulser electroporation unit (Bio-Rad Laboratories, USA) following the manufacturer’s instructions. Stock of K599 cells were refreshed for two days at 28**°**C and 225 rpm with appropriate antibiotics. About 24 hours before plant transformations a portion of the refreshed stock was transferred to a final culture volume (10x dilution) containing the same antibiotics and was left to shake at 28**°**C and 225 rpm.

The binary vector used to deliver GUSPlus (Jefferson Richard and Mayer Jorge 2003) into sunflower was pORE-E4 (Coutu et al. 2007). The GUSPlus coding sequence was extracted by restriction digestion of pANIC8E with *Sac*I and *Pst*I and was then ligated into pORE-E4 using T4 DNA ligase (Thermo Scientific) following manufacturer’s instructions. For Gateway Cloning the GUSPlus sequence was amplified with primers containing attB Gateway sites (GUSPlus-B1 5’ caagtttgtacaaaaaagcaggctatggctactactaagcatt 3’, GUSPlus-B2 5’ ccactttgtacaagaaagctggttcacacgtgatggtgatggt 3’). GUSPlus was amplified by PCR using Gateway specific primers and cloned in pDONR™/Zeo via BP clonase (Thermo Fisher Scientific). The PCR was performed following manufacturer’s instructions for Invitrogen™ Platinum™ SuperFi™ Green PCR Master Mix (Thermo Fisher Scientific). The product of the BP reaction was then transformed into SIG10-5 alpha chemically competent *E. coli* cells (Sigma-Aldrich, St. Louis). Plasmids were then extracted following manufacturer’s instructions from Thermo Scientific GeneJET Plasmid Miniprep Kit. Purified plasmids were sent for sequencing to the DNA Core Sequencing Facility, 1201 W. Gregory Drive, 334 ERML, Urbana, IL 61801. One plasmid entry clone with the desired insert confirmed by restriction digestion and sequencing was then used in an LR reaction. The LR reaction was done with two separate binary plasmids pMDC32 (Curtis and Grossniklaus 2003) and pUBQ10 (Michniewicz et al. 2015). Both vectors are Gateway® compatible and instructions outlined by Gateway® Technology were followed to complete the LR reaction and transformed in SIG10-5 alpha chemically competent cells (Sigma-Aldrich) following the manufacturer’s instructions. All binary plasmids were verified by double restriction and transformed into electrocompetent *Agrobacterium rhizogenes* strain K599. The insert was then verified once more by double restriction digestion using the same enzymes listed above.

### Seed germination and *A. rhizogenes* inoculation

Seeds were rinsed for 10-15 minutes in de-ionized water then placed in a 1.6% sodium hypochlorite solution for 20 minutes and agitated continuously. Three separate changes of autoclaved de-ionized water for 10 minutes each were used to remove all residual bleach. Clean seeds were evenly spaced in small plastic pots containing vermiculite or Pro-Mix (Premier Horticulture, Canada) potting soil and watered regularly with deionized water. Seedlings were grown under grow lights with 16 h light and 8 h dark. Just after reaching the VE stage, defined as emergence of the seedling with the first true leaves smaller than 4 cm (Schneiter and Miller 1981), seedlings were cut below the cotyledons and placed in Rockwool plugs, 2.5 cm × 2.5 cm × 4 cm tall, and soaked with 9 mL *A. rhizogenes* solution containing 1/4 MS salts (Murashige and Skoog 1962) and Gamborg’s vitamins (Gamborg et al. 1968). Explants were grown under the same conditions for two weeks and watered every other day.

### Plant cultivation after hairy root emergence

After two weeks explants were removed from Rockwool. Explants with well-formed teratomas were placed in vermiculite for an additional 10 days to allow transgenic root development. Plants were fertilized once with 5mL of full-strength MS media at the fifth day. After 10 days in vermiculite, plants were removed and stained with GUS staining solution to determine the proportion of transgenic roots present. In one trial, plants were placed in soil after removal of non-transgenic roots. These plants were grown to maturity and seeds were collected to determine the viability of the composite plants.

### β- Glucuronidase detection

To verify transgenic roots, plants’ teratomas and roots were removed from the shoots of explants and placed into GUS assay solution overnight at room temperature according to the requirements outlined in Vitha *et al*., (1995). The enzymatic reaction was stopped the following morning by replacing GUS solution with 70% ethanol. Ethanol was replaced as needed until all soluble pigments had been removed from plant material. Transgenic roots were then quantified by visual selection.

### β-Glucuronidase Quantification

GUS quantification was performed as previously described (Chen et al. 2013) on Fluoroselect flurometer (Sigma-Aldrich) according to manufacturer’s instructions. Samples were collected by cutting the teratomas with roots from 3-5 transformed plants, flash frozen in liquid nitrogen and processed with a MiniBeadBeater (Biospec Products, Bartlesville, OK). Total protein content of the samples was determined using the BIO-RAD Quick Start™ Protein Assay Kit (Hercules, CA). Protein concentration was determined based on manufacturer’s instructions. Enzyme activity was then calculated as pmole 4-MU/min/μg protein.

### Cloning of HaLBD16

RNA was extracted from leaves and roots of sunflowers from the genotype HA412-HO using the Spectrum™ Plant Total RNA Kit (Sigma-Aldrich). Single strand copy DNA (ss-cDNA) was produced from RNA using the RevertAid First Strand cDNA Synthesis (Thermo Scientific) following manufacturer instructions. The HaLBD16 was amplified by PCR using Invitrogen™ Platinum™ SuperFi™ Green PCR Master Mix (Thermo Fisher Scientific) with Gateway primers (HA LBD16-B1 5’ ggggacaagtttgtacaaaaaagcaggctatggcaactgttgctgctgg 3’, HA LBD16-B2 5’ ggggaccactttgtacaagaaagctgggtttagttcctcatcattctaac 3’). The PCR reaction was performed following manufacturer’s instructions for Invitrogen™ Platinum™ SuperFi™ Green PCR Master Mix (Thermo Fisher Scientific). The attB-flanked PCR product of LBD16 was cloned in pDONR™/Zeo, verified by sequencing, and finally cloned in binary vectors. The Gateway binary vectors for RNAi (pGMDC-RNAi, with RNAi cassette from pANIC8E) and overexpression (pGMDC32) were based on pMDC32 with GUSPlus under the tCUP promoter (from pORE-E4-GusPlus, see above). Binary plasmids clone with the desired insert confirmed by restriction digestion transformed into electrocompetent *Agrobacterium rhizogenes* strain K599. The insert was then verified once more by restriction. Composite plant production was done as described above. Five composite plant was grown in hydroponics in ¼ MS media (changed weekly) with constant aeration and 16h day photoperiod. After three weeks plant roots were stained and one main root per plant (with strong GUS staining) were scanned on Epson Pefection V850. Images were analyzed with Rhizo-II Root Biometrics Suite is a software package (Shahzad et al. 2018).

### Statistical Analysis

For statistical analysis, using IBM SPSS Statistics (IBM Corporation, Inc) or Daniel’s XL Toolbox (Kraus 2014), ANOVA was run using a Levene’s test to check for homogeneity of variance. Significance value was determined as p<0.05. Microsoft Excel 2016 was used to produce graphs.

## Introduction

The common sunflower (*Helianthus annuus L*.), is one of a small suite of crops native to North America (Putnam et al. 1990). With a decrease in export demand sunflowers have been pushed onto more marginal land where they are more likely to encounter abiotic stresses (Putnam et al. 1990) and where their production is limited (Škorić 2014). Being grown in mostly semi-arid areas, the sunflower is relatively resistant to a wide range of conditions. Sunflowers are more readily adapted to cold with seedlings often surviving freezing temperatures. Semi-arid conditions also often subject sunflowers to drought (Putnam et al. 1990). Drought and soil salinity are two major factors that influence the emergence and establishment of sunflower seedlings (Kaya et al. 2006). Different abiotic stresses such as nitrogen starvation, drought, cold, heat, and salinity reduce crop yields and the genetic potential to increase the production is not reached, indicating that the development of cultivars with an increased adaptation to environmental changing conditions should be undertaken (Miyazawa et al. 2011). Sunflower is described as normally susceptible to low temperatures and salinity, but with a relative tolerance to drought stress due to its highly explorative root system. However, available information on gene expression in response to abiotic stresses in sunflower is still limited to few works (Pradhan Mitra et al. 2014; Vitha et al. 1995)

Stable genetic transformation protocols using *Agrobacterium tumefaciens* to transform *Helianthus annuus* exist but they often present low efficiencies (Sujatha et al. 2012) and therefore are not well suited to medium-high throughput functional characterization studies (Davey and Jan 2010). For these reasons there have been very few functional studies involving transgenic *Helianthus* to date (Hewezi et al. 2006; Rousselin et al. 2002). When genes involved in root biology are under investigation, like those associated with water and nutrient uptake, *A. rhizogenes* can be used to bypass traditional labor-intensive and costly stable genetic transformation methods (Kereszt et al. 2007) by using composite plants. Composite plants are chimeric in nature, having transgenic roots and wild-type shoots, and offer a time-saving alternative to the often laborious and inefficient stable genetic transformations produced in tissue culture. (Collier et al. 2005; Taylor et al. 2006). Given efficient root transformation, composite plants are an alternative to stable transgenic lines (Estrada-Navarrete et al. 2006). Eliminating the tissue culture step, coined the *ex vitro* method, allows for a decrease in time for the production of transgenic plant material and an alternative to producing stable transgenic lines (Collier et al. 2005; Estrada-Navarrete et al. 2006). A significant reduction in plant production cost means that large scale gene characterization studies can be implemented efficiently (Collier et al. 2005; Michalec-Warzecha et al. 2016).

Functional studies have been used to characterize genes that lead to helpful discoveries for crop plants. In rice, PSTOL1 (phosphorous starvation tolerance) when over-expressed improves yield and biomass production on soil poor in phosphorous (Kochian, 2012). Genes for the increase in root depth have also been identified. The DRO1, deeper rooting 1, is a gene that when expressed in cultivars previously known for shallow root architecture increases deep rooting and overall yield by approximately ten percent (Arai-Sanoh et al., 2014). With the sunflower genome recently published (Badouin et al. 2017) there is great opportunity to perform functional studies to identify genes that may have similar impacts on productivity in the future.

## RESULTS AND DISCUSSION

### Protocol optimization

The main aim of this work was to establish an efficient system to produce composite sunflowers (Fig. 1, Supplemental Text, Supplemental Fig. 1). For development of this protocol we used Taylor *et al*. (2006) as a template and sunflower variety Peredovik. This is an older, open-pollinated variety originally developed in USSR (Arias and Rieseberg 1995). Peredovik was chosen because it has been the variety of choice across multiple fields of study (Albourie et al. 1998; Lin et al. 1975; Macías et al. 2002). The binary vector used to optimize the procedure was based on pORE-E4 (Coutu et al. 2007). This plasmid contains a plant dicot promoter tCUP with high expression at a relatively constant level thought all tissues. For the monitoring the transformation process, we introduced reporter protein GUSPlus (Broothaerts et al. 2005) to produce pORE-E4 GUSPlus binary vector. *A. rhizogenes* K599 (Mankin et al. 2007) with pORE-E4 GUSPlus binary vector was used. Clean sunflower seeds were evenly spaced in small plastic pots containing vermiculite or potting soil and watered regularly with deionized water. Seedlings were grown under 16h day. Just after reaching the VE stage, defined as emergence of the seedling with the first true leaves smaller than 4 cm (Schneiter and Miller, 1981 (Schneiter and Miller 1981), seedlings were cut below the cotyledons and placed in Rockwool plugs *A. rhizogenes* solution containing. After two weeks explants were removed from Rockwool. Plantlets with well-formed teratomas were placed in vermiculite for an additional 10 days to allow transgenic root development and GUS was used to determine the proportion of transgenic roots present.

**Fig. 1.**
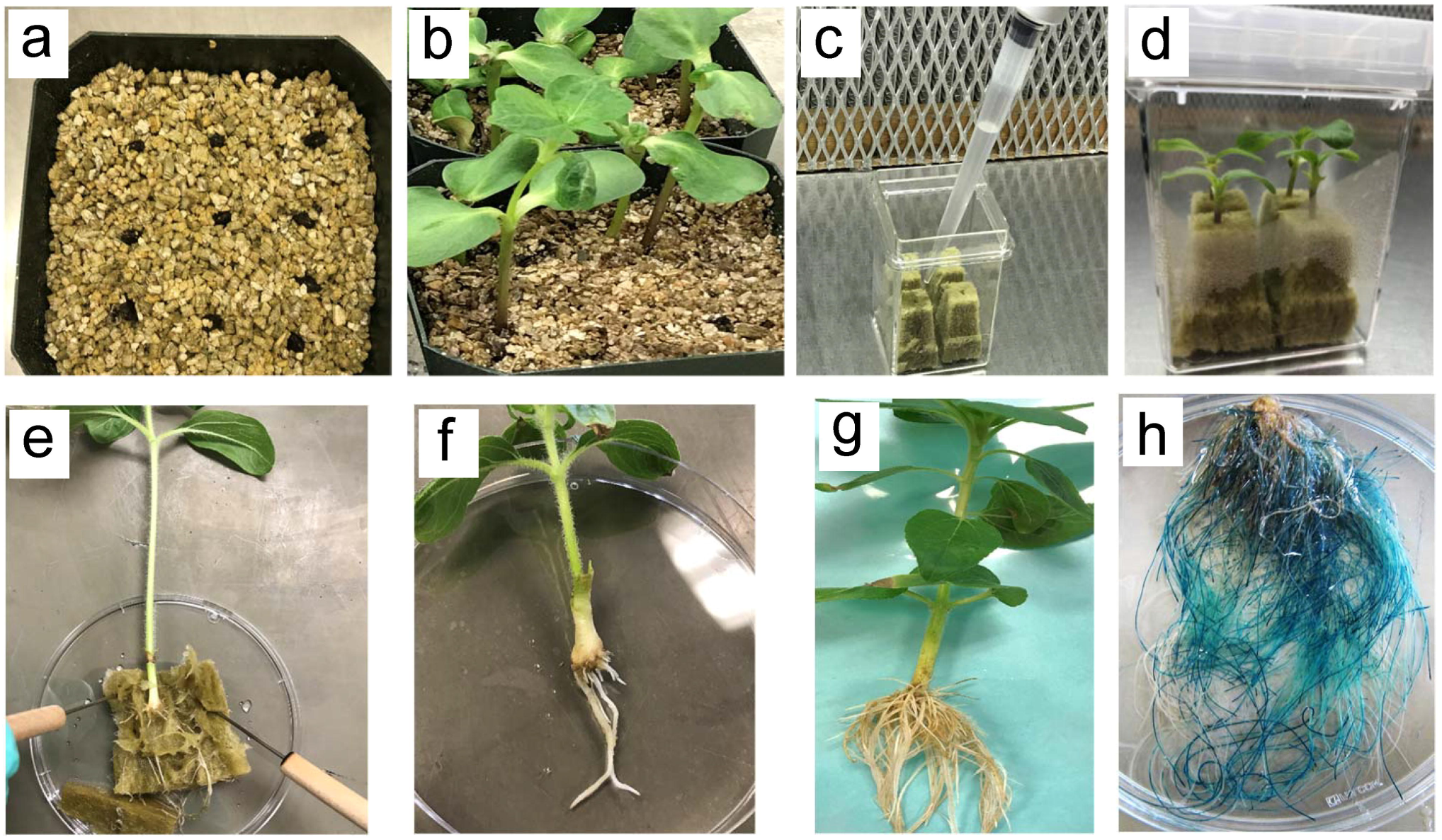
Main stages in the sunflower composite plants protocol. **a** Seeds placed just under the surface of vermiculite before being lightly covered. **b** Seedlings growing, nearing the optimum maturity for transformations about 10-14 days. **c** Rockwool being soaked with ¼ MS media and A. rhizogenes at an OD of 0.2 at 600nm. **d** Plants are placed into the pre-formed holes in rockwool from the 1ml pipet tips. **e** Composite plant removed from Rockwool after 14 days cocultivation/transformation. **f** Roots of composite plant after being removed from Rockwool. **g** Roots of composite plant after 2 additional weeks in vermiculite. **h** Roots transformed with pMDC32-GUSPlus removed from hydroponics after three weeks and stained overnight for GUS. Note, most roots are transgenic/blue.

The first change made to original the procedure was the removal of the drying step. The original protocol asserted that drying to the wilting stage was necessary for a proper transformation to occur. After trials with the drying step resulted in wilted plants that never recovered, the drying step was removed from the protocol. Instead, four Rockwool plugs were placed in Caisson boxes and sealed with vent top lids. These lids keep humidity high but allow for drying to occur at a slower pace than the wilting step. When sealed in the Caisson boxes, even with the vent-top lids, humidity reached a critical point that encouraged the growth of mold. To stop the growth of any contaminants that may have entered either on the plants themselves or by result of performing the transformation, lids were removed from the boxes after three days and watered every 48 hours until they were removed from the Rockwool. This greatly reduces the ability for fungi to spread throughout the boxes; by allowing the plugs to dry for 24 hours before being watered again most cases of fungal infection were eliminated. It was observed that sterilization increased cases of bacterial stem wilt, a disease that rarely affects sunflower seedlings, but likely persisted in small, protected areas of the hulls of the seeds. When planted un-sterilized in Pro-mix potting soil, no cases of bacterial stem wilt were observed. The original protocol (Taylor et al. 2006) outlined a bacterial optical density between 0.2-0.5 OD_600_.

At higher optical densities, 0.5 OD_600_, seedlings showed a hypersensitivity response (Kuta and Tripathi 2005). Hypersensitivity happens when the cells directly infected with *Agrobacterium* die (Sujatha et al. 2012) results in a seedling that turns black from the base and moves up the hypocotyl. An OD_600_ 0.2-0.3 provided good transformation efficiency, and no plants were lost due to hypersensitivity. Cotyledons were removed to better facilitate the placement of seedlings into Caisson boxes for transformation. It was observed that seedlings appeared healthier when cotyledons were removed and were more vigorous and less prone to disease. The average number of adventitious roots also showed a significant drop when cotyledons were removed (Fig. 2a). This makes removal from Rockwool easier and reduces the total number of roots thereby increasing the proportion of transgenic roots.

**Fig. 2.**
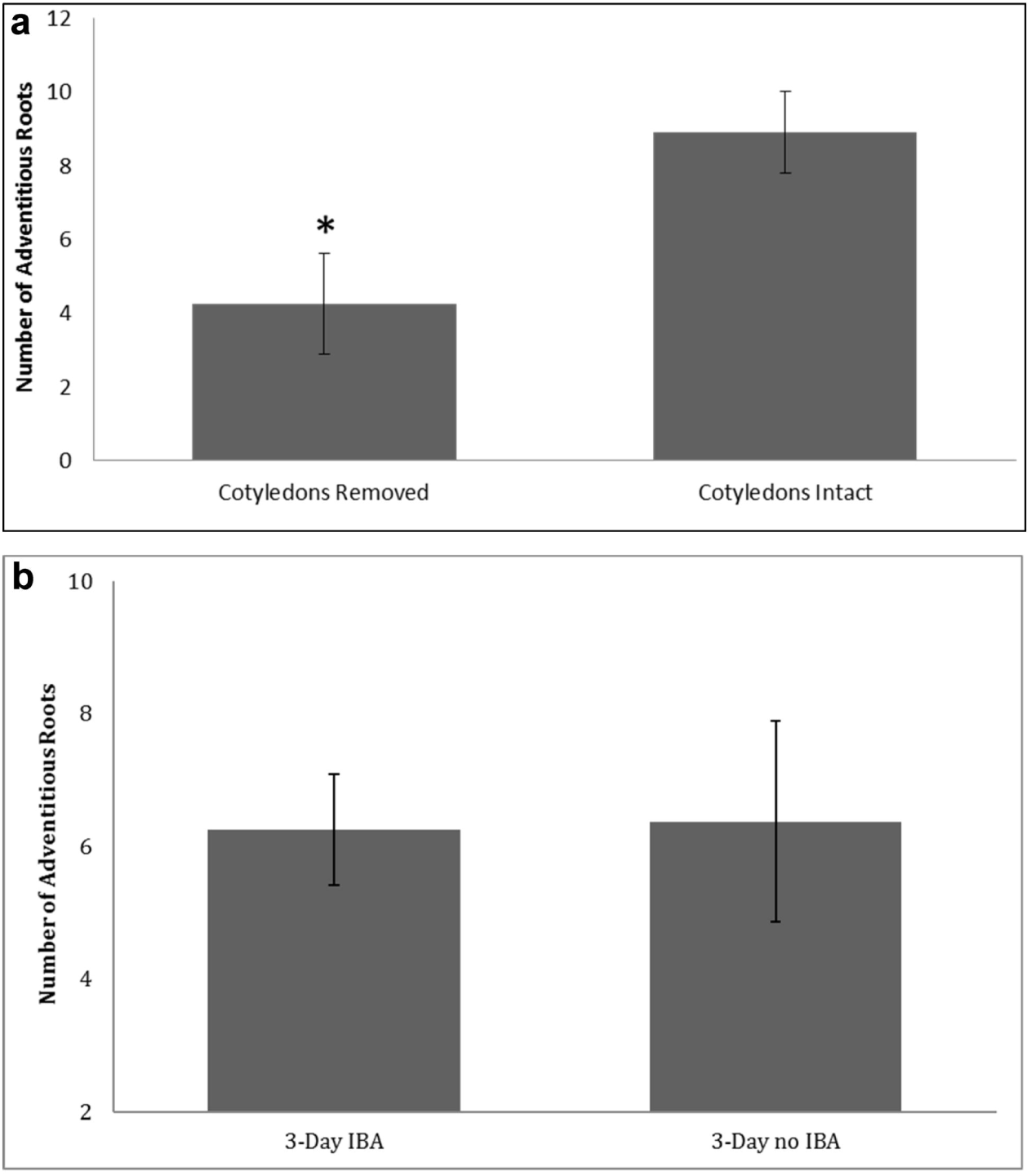
Effect of cotyledons and IBA on roots development. **a** Effect of cotyledons presence on root formation. Average number of adventitious wild-type roots showing a significant decrease when cotyledons were removed * p<0.05. All plants were grown under 16 h light and 8 h dark for the duration of the experiment. Values are averages of 10-12 plants ±SE. **b** Effect of IBA on root formation. Average number of adventitious roots showing non-significant differences between a treatment that moved explants to new Rockwool plugs treated with IBA after 3 days in co-cultivation and the treatment that moved explants to plugs with only ¼ MS media after three days of co-cultivation with *A. rhizogenes*. Values are averages of 8 explants ± SE.

Many protocols advise using the stem tissue of mature plants (Collier et al. 2005; Taylor et al. 2006). It was found that sunflower transformations provided better efficiency using hypocotyls (Everett et al. 1987). In our protocol, when using hypocotyls at least 80% out of the plants readily formed teratomas. When using mature stem tissue, no plants formed teratomas. Hypocotyls were determined to be the most responsive tissue for transformations (Benzle et al. 2015). Sunflowers are also determinant in their flowering (Sujatha et al. 2012) so it is critical to start the transformation process as quickly as possible. Auxins, like IBA (indole 3-butyric acid), are often helpful in the formation of roots and can increase transgenic root formation (Li and Leung 2003). In our case, IBA had a negligible effect on the formation of transgenic roots but did increase the number of non-transgenic adventitious roots the average plant formed. IBA provided no increase in transgenic roots and increased the proportion of non-transgenic roots so the treatment with IBA was not used (**Fig. 2b**). Transgenic root development was slower in this initial phase than the development of adventitious roots as the development of a teratoma is required before roots will start to grow. The removal of nearly all the roots developed during the two weeks in the Rockwool allowed the transgenic roots to developed when placed in the vermiculite. Explants are then removed from vermiculite, washed and can further grow in soil or hydroponics. Several composite plants were grown to maturity in soil in 4” trade pots. These plants flowered and produced seeds (Fig. 3), and had about 12 seeds/plant, with high viability (96%).

**Fig. 3.**
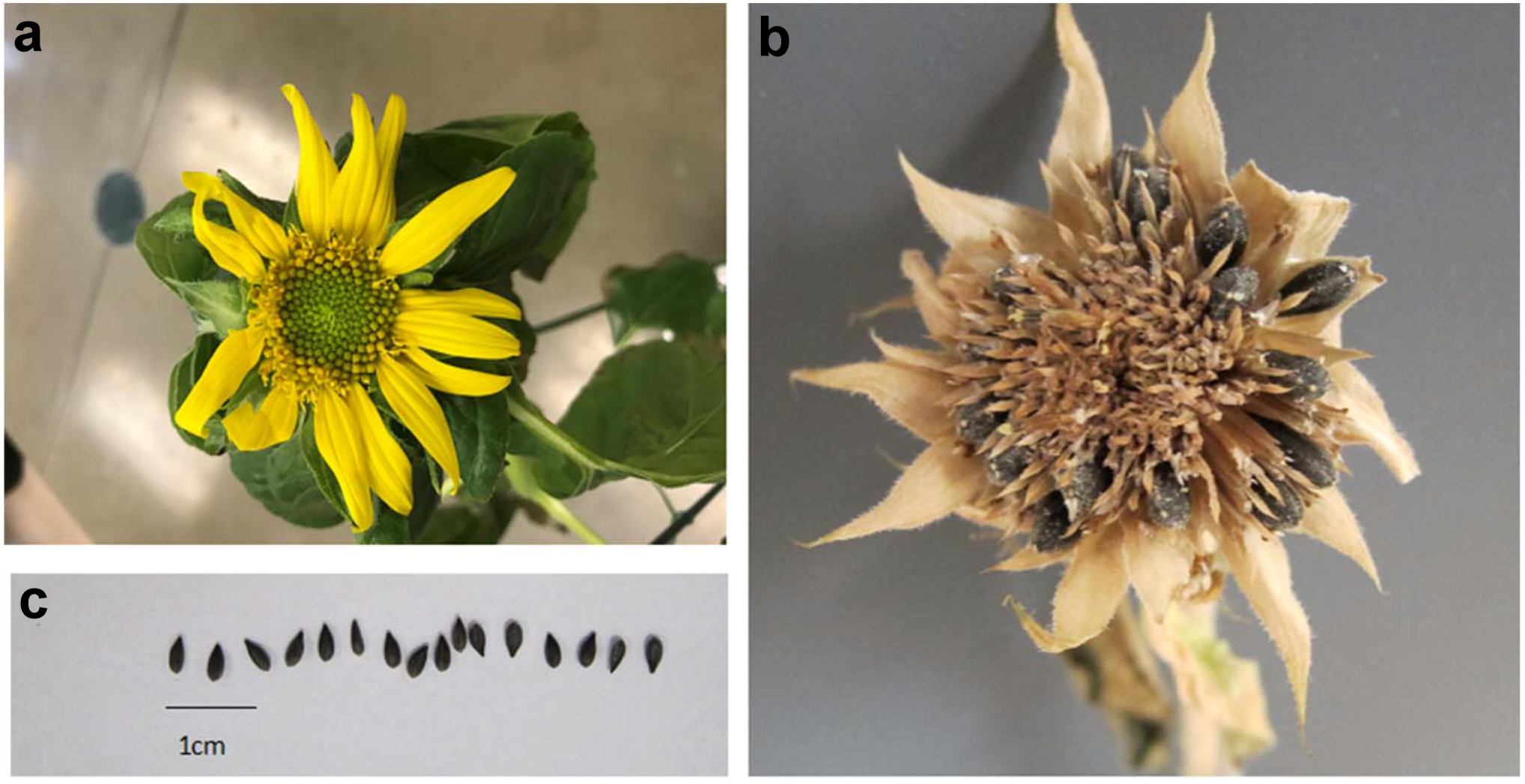
Complete development of composite plant grown in soil. **a** Fertile flower starting pollination. **b** Dry flower head with seeds around the periphery. **c** Seeds from composite plants.

### Variety testing

Multiple varieties were tested for transformation efficiency (Fig. 4). Out of thirteen varieties tested three groups were apparent: those with good transformation efficiency (Peredovik, HA280, HA298, Mammoth, HA311, and RHA801), those with medium efficiency (RHA271, HA412, HA115), and those with poor efficiency (HA89, HIR34, HA412-HO, and HA236). Genotype specificity is not uncommon in transformations (Han et al. 2000; Heeres et al. 2002; Hoffmann et al. 1997; Lee et al. 2004; Zhang and Finer 2016). Similarly, in another study, that used cotyledons of mature seeds, sunflowers had variable transformation frequency (Sujatha et al. 2012). For the genotypes that were shown to be recalcitrant it may be possible to improve the efficiency by using acetosyringone, which has been shown to induce the *vir* genes of *Agrobacterium* and increase the transformation rate (Guivarc’h et al. 1993).

**Fig. 4.**
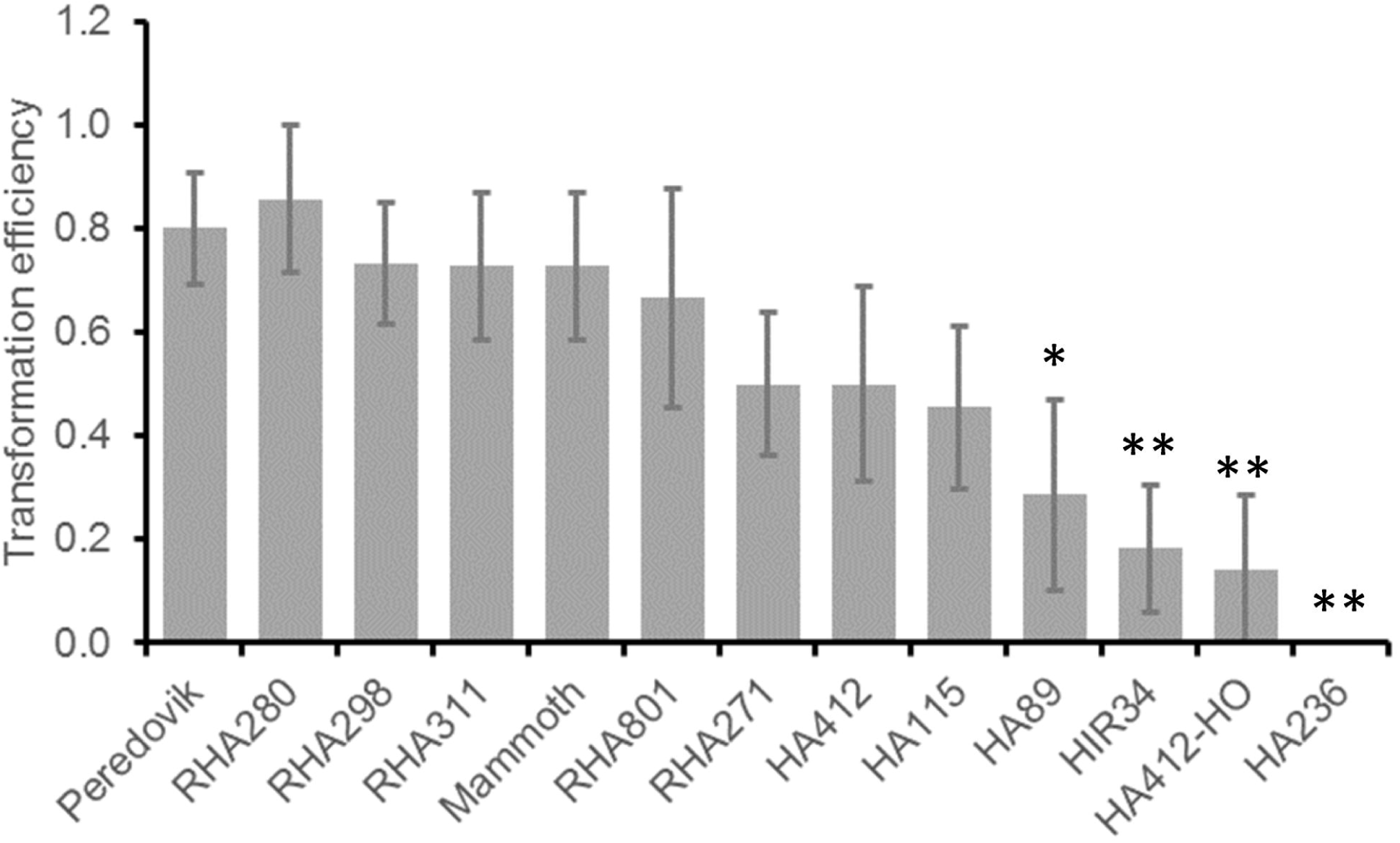
Transformation efficiency ratio of various sunflower varieties. Proportion of plants that formed at least one transgenic root. Significantly different proportions denoted by * one * denotes p<0.05, two ** denotes significance level of p<0.01.

### Constitutive promoters testing

We tested two more constitutive promotes for efficiency in roots expression, double CaMV35S (2xCaMV35S) and ubiquitin (from Arabidopsis) promoters in plasmids pMDC32 (Curtis and Grossniklaus 2003) and pUbi10 (Michniewicz et al. 2015), respectively (Fig. 5). The 2xCaMV35S promoter was expressed the most highly out of three tested promoters. Promoters tCUP2 and 2xCaMV35S were expressed throughout the roots, while ubiquitin appeared to be expressed more in the vascular tissue of the root. Promoter activity of tCUP2 and CaMV35S have been reported to be expressed at similar levels in other studies (Coutu et al. 2007). The ubiquitin promoter is typically reported to have more moderate expression levels in most plants (Geldner et al. 2009). These results demonstrated that all three promoters tested are useful for sunflower transformation.

**Fig. 5.**
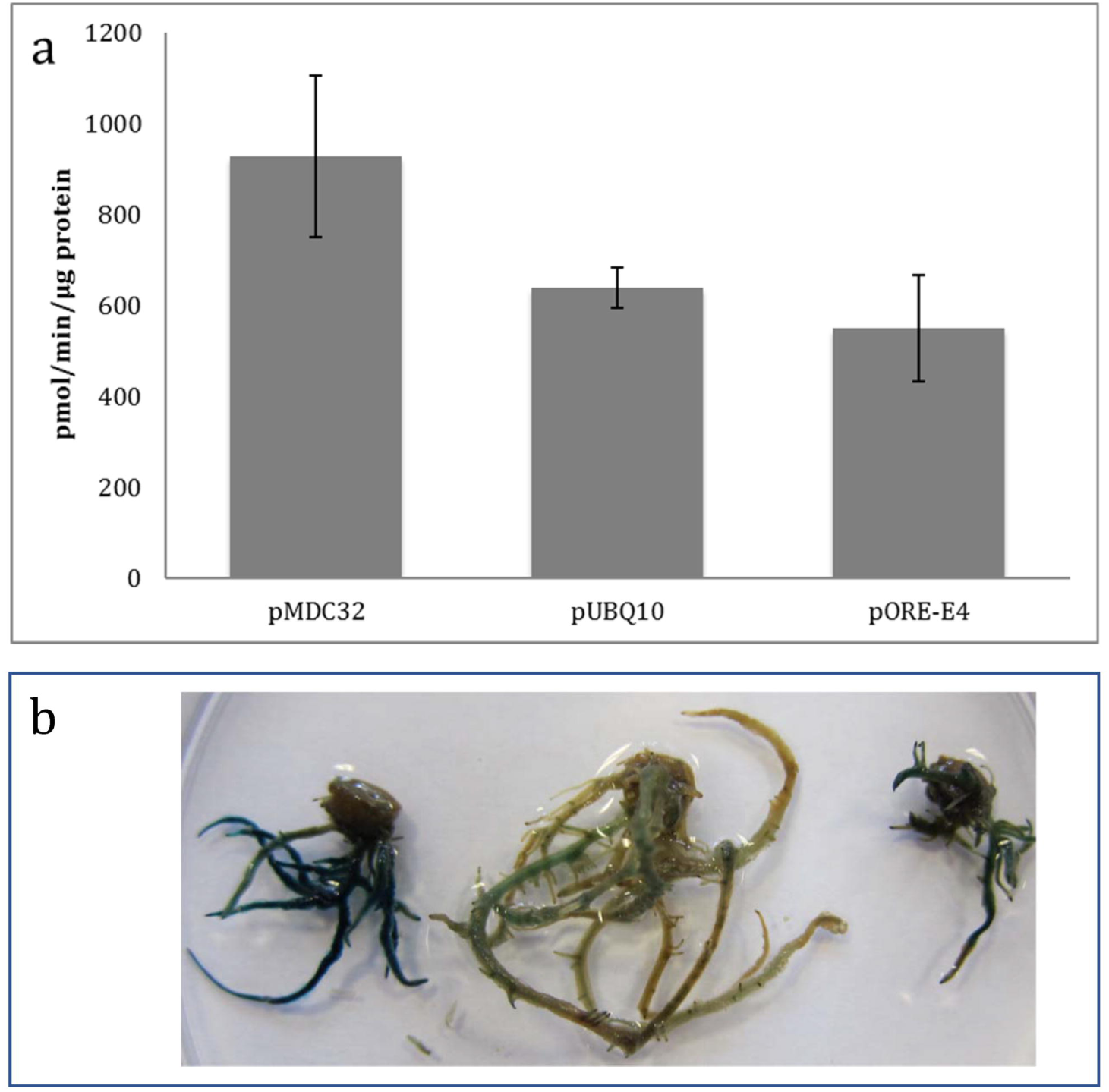
Activity of the three constitutive promoters. **a** Promoter activity based on fluorometric GUS analysis. Values are averages of ∼3 plants ± SE. **b** Roots stained for GUS activity. Note, differential expression is visualized between pMDC32, pUBQ10, and pORE-E4, respectively.

### HaLBD16 transgenics characterization

Genes in the Lateral Organ Boundaries Domain (LBD) family have been shown to influence the development of lateral root primordia (Lee et al. 2009; Okushima et al. 2007). Previous studies involving *Arabidopsis thaliana* mutants lacking the LBD16 showed a reduction in lateral root formation, while transformants with an overexpression cassette containing LBD16 showed a significant increase in lateral root production (Feng et al. 2012). LBD16 is important in the auxin response of lateral root formation (Fan et al. 2012; Lee et al. 2009). Here we conduct a functional study to characterizing the sunflower LBD16 homolog (HaLBD16, gene ID HanXRQChr10g0281141, protein ID XP_021984895.1) with both an over-expression and down-regulation gene constructs. Plant roots transformed with LBD16 over-expression (16OE) and LBD16 RNAi (16i) were produced using the method described above. As previously reported for the effect of the LBD16 in Arabidopsis (Feng et al. 2012), the sunflower roots of the 16OE plants showed more lateral roots (branching) than 16i roots (Fig. 6). Moreover, the same gene was identified in the region involved in root biomass in sunflower (Masalia et al. 2018). This result demonstrate that the composite plants approach can be successfully used for functional gene studies in sunflower.

**Fig. 6.**
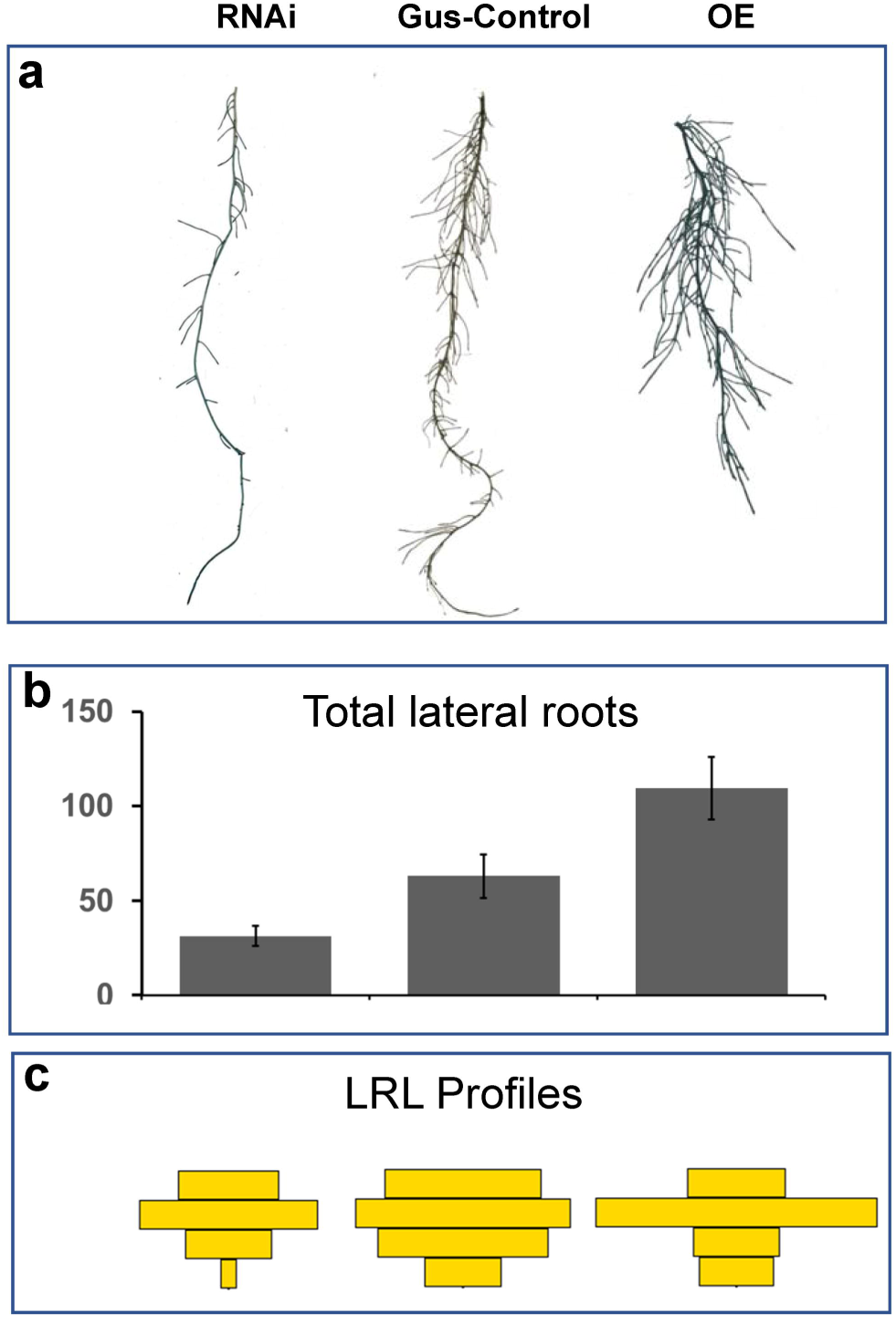
Expresssion of HaLBD16 transcription factor affects roots architecture. **a** Typical transgenic root from plants transformed with the three constructs: HaLBD16-RNAi, Control pORE-E4-GusPlus, HaLBD16-OE, respectively. **b** Expression of the HaLBD16 increase total lateral roots n=5 ± SE. **c** Lateral roots profiles, represented as the main root was divided into four sectors and the mean lateral roots lengths within each sector is represented as a rectangle over the sector n=5.

Composite plants are a fast and efficient way to perform characterization studies in roots of many non-model plants. With the availability of the sunflower genome there is a need for a system in which to perform functional studies efficiently. A cost effective and time efficient way to generate transgenic tissue for studies is with composite plants via inoculation with *A. rhizogenes*. With the protocol developed here functional characterizations can be carried out in sunflower with efficiency and in a short time span (∼2 months). This has the potential to characterize many new candidate genes regulating root development and stress adaptation.

## Acknowledgements

We would like to thank Prof. Gary Stacey (University of Missouri) for providing the K599 strain, and the Department of Biological Sciences for the funding: startup found (YSY), Redden grants (YSY), and Lewis Hanford Tiffany Botany Graduate Research Fund (TP).

## Author contributions

TP and YSY designed the experiment, collected and analyzed data, and drafted the manuscript. TP conducted the research and collected the data.

## Conflict of interest

The authors declare they have no conflicts of interest.

