## Supplemental Text for "Composite plants for a composite plant: An efficient protocol for root studies in the sunflower using composite plants approach"

### Step-by-step protocol for the composite plants production in sunflower

Planting seeds

- Using deionized water and potting soil prepare a seedbed.
- After the soil is lightly packed into a tray gently press the sunflower seeds in until they are just below the surface.
    - It is helpful to face seeds radicle down to improve germination and uniformity among seedlings.
- Cover lightly with a fine layer of soil, and water carefully so as not to uncover the newly planted seeds.
- Place seed trays under a grow light for 1.5 to 2 weeks at 25±3°C, 16h day.

*A. rhizogenes* preparation

- Estimate three days before the first set of true leaves reach 4 cm in length, refresh the *A. rhizogenes* stock culture and let it incubate at 28°C at 225 rpm for 2 days.

- 1 day before seedlings reach ideal size use 1mL of stock culture for every 10 mL of final culture required for the trial.

  10 mL of final culture volume will normally cover about 10 plants.

- Incubate at the same temperature and rpm for 24 hours.

Transformations

- Sterilize Caisson boxes with vent top lids and Rockwool plugs already in place. It is easiest to press holes into Rockwool before sterilizing using a clean 1 mL pipet tip.

- Spin down the bacteria at 3300g for 10 min at room temperature and re-suspend in ¼ MS with vitamins.
- Under a sterile hood, dilute the bacteria to an OD of 0.2 at 600 nm using more of the ¼ MS

  Typically, a 13x dilution will suffice.

- After the proper density is achieved apply 9 mL of the bacterial solution to each Rockwool plug that has already had a hole pressed into it.
- Using scissors remove cotyledons and cut the seedling 2-4 cm from the base of the first true leaves. Using forceps gently press the seedling into the Rockwool taking care to not pierce the hypocotyl.
- After all 4 individuals are in the Caisson box cap with a vent top lid and set them under a grow light for 3 days undisturbed.
- After 3 days, to reduce humidity and threat of disease it is best to uncap the boxes and set the lids ajar on top of the boxes.
- Water every other day or as the plugs begin to dry.

Removal from Rockwool

- After 14 days remove the Rockwool from the sunflowers. This can be achieved with two dissecting needles.

- Note: Removal is easier if the Caisson boxes are flooded with water for 6-8 hours before removal, this softens the Rockwool and allows roots to be removed undamaged.

- Remove obvious non-transgenic adventitious roots. Anything of great size or not originating from the teratoma is unlikely to be transgenic. Likewise, transgenic roots can be identified using markers, such as GUS/GFP/RFP.
- Place the sunflowers into a Caisson box with vermiculite and keep them watered for 2 weeks.
- After 5 days in vermiculite fertilize each Caisson box with 5mL of full-strength MS media to produce more vigorous plants.
- After 14 days well developed plants can be transferred in soil or hydroponics. Transgenic roots can be identified using visual markers, such as GUS/GFP/RFP.
