## Supplemental Fig. 1 for "Composite plants for a composite plant: An efficient protocol for root studies in the sunflower using composite plants approach"

**Supplemental Fig. 1.** Timeline for composite sunflower production. Pictures are the same as in the Fig.1 and are used to envision the main outcome for the stages in the procedure.

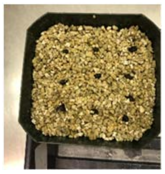

### Day 0

Plant seeds

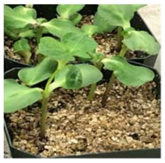

### Day 1 - 14

Grow plantlets

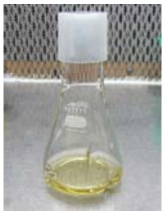

### Day 12

Culture bacteria

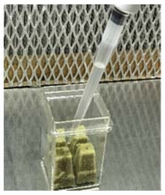

### Day 15

Rockwool plug with *A. rhizogenes*

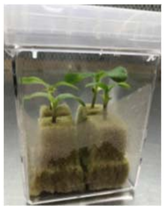

### Day 16 - 29

Plant transformation

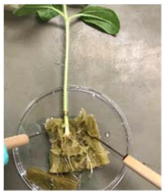

### Day 30

Removal from rockwool

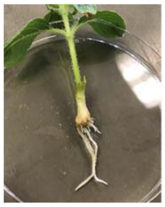

### Day 31 - 44

Cultivation in vermiculite

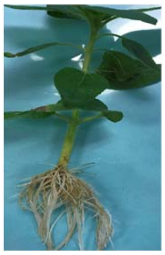

### Day 45

Composite plants transfer to soil/hydroponics

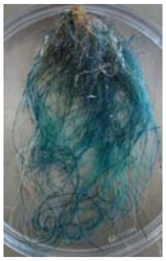

### Day 65

Characterization of the fully developed roots
